# PENet: A phenotype encoding network for automatic extraction and representation of morphological discriminative features

**DOI:** 10.1101/2023.05.21.541653

**Authors:** Zhengyu Zhao, Yuanyuan Lu, Yijie Tong, Xin Chen, Ming Bai

## Abstract

Discriminative traits are important in biodiversity and macroevolution, but extracting and representing these features from huge natural history collections using traditional methods can be challenging and time-consuming. To fully utilize the collections and their associated metadata, it is urgent now to increase the efficiency of automatic feature extraction and sample retrieval. We developed a Phenotype Encoding Network (PENet), a deep learning-based model that combines hashing methods to automatically extract and encode discriminative features into hash codes. We tested the performance of PENet on six datasets, including a newly constructed beetle dataset with six subfamilies and 6566 images, which covers more than 60% of the genera in the family Scarabaeidae. PENet showed excellent performance in feature extraction and image retrieval. Two visualization methods, t-SNE, and Grad-CAM, were used to evaluate the representation ability of the hash codes. Further, by using the hash codes generated from PENet, a phenetic distance tree was constructed based on the beetle dataset. The result indicated the hash codes could reveal the phenetic distances and relationships among categories to a certain extent. PENet provides an automatic way to extract and represent morphological discriminative features with higher efficiency, and the generated hash codes serve as a low-dimensional carrier of discriminative features and phenotypic distance information, allowing for broader applications in systematics and ecology.

## Introduction

Discriminative traits are of particular importance in biodiversity and macroevolution, as they provide crucial information for species delimitation, systematics relationship assessment, and phenotypic evolutionary analysis (Ericson, 1997; Koehl, 1996; Wiens, 2001; Wiens & Servedio, 2000). Traditional methods to extract discriminative phenotypic traits are typically relatively subjective and rely on experiential expertise (Hawkins, 2014). Manual input is still required, even with quantitative methods available, such as morphometrics and geometric morphometrics (Ibacache et al., 2010; Rohlf & Marcus, 1993). Besides extracting discriminative features, further utilizing these features to search for similar phenotypic individuals in a diverse range of natural resources can pose a big challenge for researchers, let alone non-experts. For example, searching for desired specimens with certain discriminative phenotypes in natural history museums can be challenging.

Natural history collections and their associated metadata (e.g., dates, locations, climate) offer a valuable resource for researchers to undertake detailed analyses and address complex questions pertaining to ecology and evolution (Lister, 2011; Winker, 2004). However, a large percentage of specimens remain uncategorized and underutilized, hindering their full potential. To address this issue, recent efforts have focused on digitizing specimens, resulting in a vast collection of digital resources, including images, 3D scans, measurements, and more (Hedrick et al., 2020; Nelson & Ellis, 2019; Page et al., 2015). Therefore, on this basis, diversification of indexing methods can assist researchers in efficiently retrieving desired specimens from natural history collections. Feature-based searching is a promising method, particularly when combined with machine learning, whereby a database of digital features is constructed and computer algorithms are used to match these features, enabling researchers to effectively retrieve desired specimens from natural history collections (Bustos et al., 2005; Tagare et al., 1997; Vishraj et al., 2022).

Recently, the rapid development of machine learning and deep learning has led to the emergence of many effective feature extraction algorithms for biological features, enabling tasks such as species classification and feature segmentation (Christin et al., 2019; Høye et al., 2021; Xiong et al., 2021). However, the extracted feature vectors from digitized collections can be highly dimensional, which presents challenges for direct utilization in specimen retrieval. In computer science, hashing methods are commonly employed to handle complex, high-dimensional data and vectors by reducing their dimensionality to hash codes, while still preserving important information (Chi & Zhu, 2018; Knott, 1975). And the hashing methods make processing and analysis more efficient, especially in tasks such as sample retrieval (Jinhui Tang et al., 2015). The hash code is composed of a certain length of 0/1 digits, for example, “11010011101011”, where “1” can be regarded as representing a certain characteristic present in the image, and “0” represents their absence. As a result, combining deep learning models as feature extractors with hash codes as feature representations has the potential for faster retrieval of sample images (Luo et al., 2020).

In this study, we propose an end-to-end phenotype encoding network with the backbone of the latest deep learning architecture Swin transformer (Liu et al., 2021), which can automatically extract high-dimensional features from input images and convert them into hash codes. We here have applied six datasets to explore the application of hash codes in two aspects. First, we verified the ability of hash codes to retrieve specimens at a large scale in scenarios such as the natural history collections in six datasets (Beetle, Fungi, Butterfly, Flower, Bird, and Fused datasets), and demonstrated the application cases of using hash codes to retrieve specimens in simulated database. Next, to further explore the representation ability of hash codes, we demonstrate the representation ability of the hash code as a whole and each bit of the hash code using two visualization methods, respectively, indicating that hash codes are excellent carriers of features. Additionally, when converting discriminative features within the images into hash codes, we effectively obtain the morphological distance matrix of these features. Therefore, we used the beetle dataset, which covers more than 60% of the genera in the six major subfamilies of Scarabaeidae, as an example to further investigate the application of hash codes generated by PENet.

## MATERIALS AND METHODS

### 2.1 Datasets and data preprocess

#### Beetle dataset

This dataset contains 6566 images (Zhao et al., 2023), all of which are the dorsal views of the beetles in the family Scarabaeidae. Specifically, it consists of six subfamilies under the Scarabaeidae, including Aphodiinae (703 images), Cetoniinae (1660 images), Dynastinae (404 images), Melolonthinae (1235 images), Rutelinae (1167 images), and Scarabaeinae (1397 images). Additionally, this data set contains more than 60% of the genera (Total ∼2175 genera) in these six subfamilies (https://www.catalogueoflife.org/?taxonKey=6278C). The images were collected from a variety of sources, including photographs taken in major museum collections around the world and images published in monographs and literature (Table S1). To ensure the reliability of the data, all images were confirmed at the subfamily level via taxonomists; on this basis, most images were identified at the species level. Thus, the beetle dataset was used to test the performance of the PENet model, while also being employed to explore the application of hash codes to systematics.

#### Fungi dataset

This dataset is derived from the Danish Fungi 2020 (Picek et al., 2022), which contains a total of 295,938 images and 1604 species, from which we selected 20 species as experimental data. Most of the images in this dataset are wild fungi, with complex backgrounds that contain not only fungi but also other elements. This dataset was used to test the performance of our model in handling wild data.

#### Butterfly dataset

The Butterflies dataset is a publicly available dataset from the Web (Gerald, 2022b). It consists of 75 species with 10,035 images.

#### Flower dataset

The flower dataset is a publicly available dataset for computer vision-related tasks that was released by Oxford University in 2008 (Nilsback & Zisserman, 2008). It contains a total of 102 species.

#### Bird dataset

This dataset includes 400 species with a total of 62,388 images(Gerald, 2022a). It comes from the same source as the butterfly dataset, but with a larger number of species and can carry out further validation.

#### Fused dataset

To validate the performance of the PENet in dealing with more complex multielement datasets, we fuse the five datasets into a fused dataset.

To ensure that the images from various sources are suitable for model training, we preprocessed the images through several steps. The images were first resized to 224×224. And during the resizing process, a solid color filling strategy was used to prevent image deformation and ensure that all images have the same length and width. Then, the datasets were divided in a ratio of 7:2:1, which means that 70% of the data is used for training, 20% for validation, and 10% for testing. This division ensures that the model is trained on a sufficiently large amount of data while also having enough data for validation and testing to assess its performance.

### 2.2 PENet and model training

In this study, we propose a new network, called the phenotype encoding network (PENet), developed through deep learning combined with hash codes, which can represent extracted features by a series of binary numbers (Figure 1a). We chose the Swin transformer (Liu et al., 2021) as the basic backbone of the network.

**FIGURE 1.**
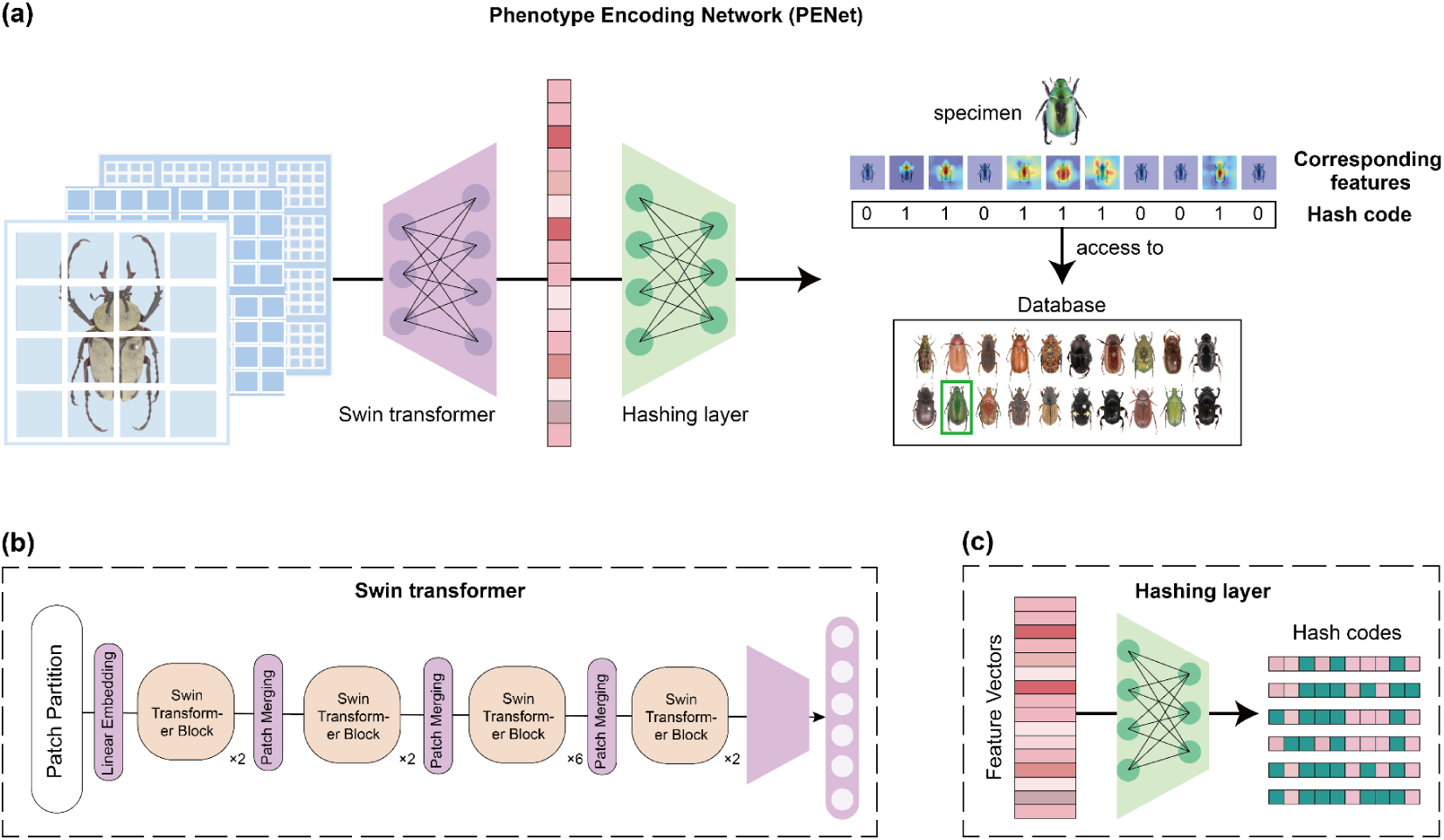
Illustrations of the PENet. (a) The PENet pipeline. (b) The architecture of the Swin transformer. (c) Hashing layer map feature vectors to hash codes.

Currently, the Swin transformer is considered a state-of-the-art deep learning model, and its architecture is distinctive compared to those of other transformer-based models in the field of computer vision (Dosovitskiy et al., 2021; Han et al., 2023). The Swin transformer adopts a hierarchical design, similar to convolutional neural networks (CNNs), with the deepening of the network layers, the receptive field of nodes is also constantly expanding. As illustrated in figure. 1b, the Swin transformer consisted of four stages, each including a Patch Merging operation (except for the first stage, which was a linear layer) and multiple Swin transformer blocks. The role of the Patch Merging module is to reduce the resolution of the input feature graph by downsampling at the beginning of each stage. And after each stage, the resolution becomes half, and the number of channels doubles. The Swin transformer block in each stage is mainly composed of two Window Attention modules, which adopt the Window based Multi-head Self Attention (W-MSA) method and the Shifted Window based Multi-head Self Attention (SW-MSA) method, respectively. And these two methods can reduce the computational complexity and take into account the association between windows.

In PENet, the Swin transformer is used to perform feature extraction on the input images. Specifically, we adjust the input dimension of the model to 224×224×3, where 224×224 is the length and width of the input image, and three is the number of channels. First, each input image was divided into 56×56 patches, where each patch is 4×4, ensuring that there is no intersection between patches. Second, embedding was performed on each patch to encode it into a 96-dimensional vector. These generated vectors were subjected to linear treatment and then successively input into Swin transformer blocks for feature extraction. Third, the extracted features were passed through the global average pooling layer to generate a 768×1-dimensional vector containing the high-dimensional features of the input image. Finally, we added a hash layer at the end of the Swin transformer to map the extracted feature vectors to hash codes of variable length, and the length of hash codes can be set. In summary, the PENet is an end-to-end model that takes images as input and produces binary hash codes as output.

During model training, adaptive moment estimation with weight decay (AdamW) was selected as the optimizer (Loshchilov & Hutter, 2019), and the loss function proposed by Liu et al. was selected (Liu et al., 2016). The central concept of this loss function is to encourage similar images to have similar hash codes and dissimilar images to have different hash codes. Based on this loss function, the model is trained by randomly selecting pairs of images as input. If two images have similar features, their hash codes keep close to each other; otherwise, they are pushed far away. This approach ensures that the model can learn the similarities and differences between different data, and accurately map high-dimensional features to the hash codes.

### 2.3 Validation of the extraction capability of the Swin transformer

In computer vision models, accuracy in classification tasks is an intuitive measure of their feature extraction capabilities. The formula for accuracy is as follows:

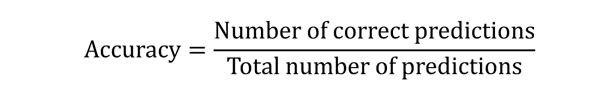

Therefore, to evaluate the feature extraction ability of the Swin transformer, we tested its classification accuracy on two datasets with significantly different categories: beetle dataset (6 categories) and bird dataset (400 categories). In this case, the parameters were optimized in the training set and the accuracy was calculated in the validation set. We also compared its performance with the two most representative convolutional neural networks, AlexNet (Krizhevsky et al., 2017) and ResNet (He et al., 2016). During the training process, all three models were trained with the same configuration. To further improve the speed of model convergence, we loaded weights that were pre-trained on ImageNet and trained for 50 epochs with a batch size of 64. And we use data augmentation strategies including random rotation, random flipping, and random center cropping in the training process. Additionally, the confusion matrix is a commonly used tool for evaluating the performance of classification models (Sokolova & Lapalme, 2009), as it provides insight into the model’s classification performance across different categories. Therefore, we performed a confusion matrix analysis on the test set that had not been involved in the training process, to further clarify the ability of different models to differentiate each category in the dataset.

### 2.4 Fast retrieval of specimen images

To demonstrate the ability of hash codes to retrieve specimens, we tested six datasets (Beetle, Fungi, Butterfly, Flower, Bird, and Fused dataset) using hash codes generated by PENet. The training set was used to adjust the model weight parameters, while the test set and the validation set were used to evaluate the performance of the model. We did this by computing the Hamming distance between the test set hash codes and the validation set hash codes. Here, the Hamming distance indicated the number of different characters in the corresponding positions of two equal strings, that is, the number of different bits in the two hash codes (Bookstein et al., 2002). The Hamming distance between two hash codes, x and y, is denoted as:

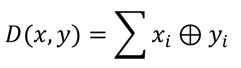

In this formula, *i* = 0, 1, …, n - 1, *x* and *y* are hash codes, n is the length of the hash code, and ⊕ denotes the exclusive or (XOR) operation.

To further evaluate the model’s retrieval capability, we used the mean Average Precision (mAP) as a metric, which measures the quality of retrieval results and the accuracy of ranking (Luo et al., 2020). The mAP is calculated as the average of the average precision of each query over all queries, which is calculated as:

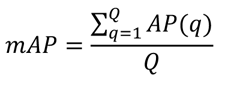

where *Q* is the number of queries, and the Average Precision is calculated as:

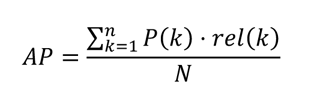

where: *n* is the total number of retrieved items, *P*(*k*) is the precision at rank *k*, *rel*(*k*) is an indicator function that is 1 if the item at rank *k* is relevant, and 0 otherwise, *N* is the total number of relevant items in the dataset.

Specifically, for each dataset, every hash code generated from the test set served as a query. During the retrieval, the query hash code was compared with every hash code in the validation set, and the Hamming distance was calculated for each comparison. The images in the validation set were then sorted according to their distance from the query, with the most similar images appearing at the top of the ranking. The mean average precision (mAP) was then calculated based on the sorted results. The size of the test set is equivalent to the number of queries. To achieve optimal performance for various hash-code lengths, we trained PENet each time the hash-code length was modified. The training was carried out for 45 epochs, and we saved the model parameters that achieved the best performance for each hash code length.

To further demonstrate the effectiveness of hash codes in retrieval and storage, we first selected 1000 images from the fused dataset that were not used in the training process to serve as a simulated database and converted them into hash codes via the PENet. Next, we selected one untrained image from each of the five datasets (Beetle, Fungi, Butterfly, Flower, and Bird datasets) and converted them into hash codes. Finally, the hash code generated in the previous step was used as a query to retrieve the simulated database and return the five most similar images among them.

### 2.5 Verification and visualization of hash codes representation capability

To demonstrate the overall representation capability of the hash codes, we applied the t-distribution stochastic neighbor embedding algorithm (t-SNE) to visualize the generated hash codes, which is currently a more general method for downscaling and visualization in the field of machine learning (Laurens & Hinton, 2008). The t-SNE algorithm is a non-linear dimensionality reduction method that can reduce high-dimensional data points to two or three dimensions while preserving the original data structure. We have chosen to use 64-bit hash codes for t-SNE visualization, with each hash code representing a 64-dimensional feature vector. For datasets other than the beetle dataset, that contain more categories, we randomly select 10 categories as samples.

The feature extraction process in deep learning has long been considered to have relatively low interpretability. As a result, some methods have emerged to attempt to visualize the extracted features, among which the gradient-weighted class activation mapping (Grad-CAM) algorithm are more commonly used (Selvaraju et al., 2020). Grad-CAM generates heatmaps by calculating feature map gradients and weighting them with average pooling, it can help to understand which features are utilized by the network. Here, a total of 30 species of beetles in six subfamilies are selected from the beetle dataset as test data for illustration. We used PENet to convert these beetle images into 64-bit hash codes and generated heatmaps using the Grad-CAM algorithm to display each bit of the hash code features.

### 2.6 Constructing phenetic distance tree based on hash codes

As the hash codes contain the discriminative features extracted by the model and the distance of morphological differences among different categories, the generated hash codes from different taxa could form a morphological distance matrix. Here, we selected two species from each subfamily of the beetle dataset to generate 64-bit hash codes using PENet (Table S2). Based on these hash codes, we constructed a phenetic distance tree, with one species of Staphylinidae selected as the outgroup. The hash code of the outgroup was also generated using PENet. These selected species did not participate in the PENet training process to avoid overfitting. We used Nona (Goloboff, 1993) run via Winclada (Nixon, 1999) to perform heuristic searches to find the most parsimonious trees (MPT). MPTs were found with a heuristic search using the commands “hold1000000”, “mult*1000”, “hold/10”, “mult*max*”.

## Result

### 3.1 Identification performance

#### 3.1.1 Beetle dataset

After 50 rounds of training, all three models, namely AlexNet, ResNet, and Swin transformer, gradually converged, with respective training times of 101, 123, and 126 minutes (Figure S1). On the validation set of the beetle dataset, their highest accuracy was 90.58%, 94.98%, and 97.8%, respectively (Figure 2a). The confusion matrix analysis revealed that the Swin transformer outperformed the other two models in the performance of each subfamily, with accuracy above 95% for every subfamily (Figure 2b). However, the other two models mainly had prediction errors concentrated in the subfamilies Dynastinae, Melolonthinae, and Rutelinae. In particular, their prediction accuracy for the subfamily Dynastinae was relatively poor, with AlexNet at 78% and ResNet at 90%, compared to the accuracy of other subfamilies.

**FIGURE 2.**
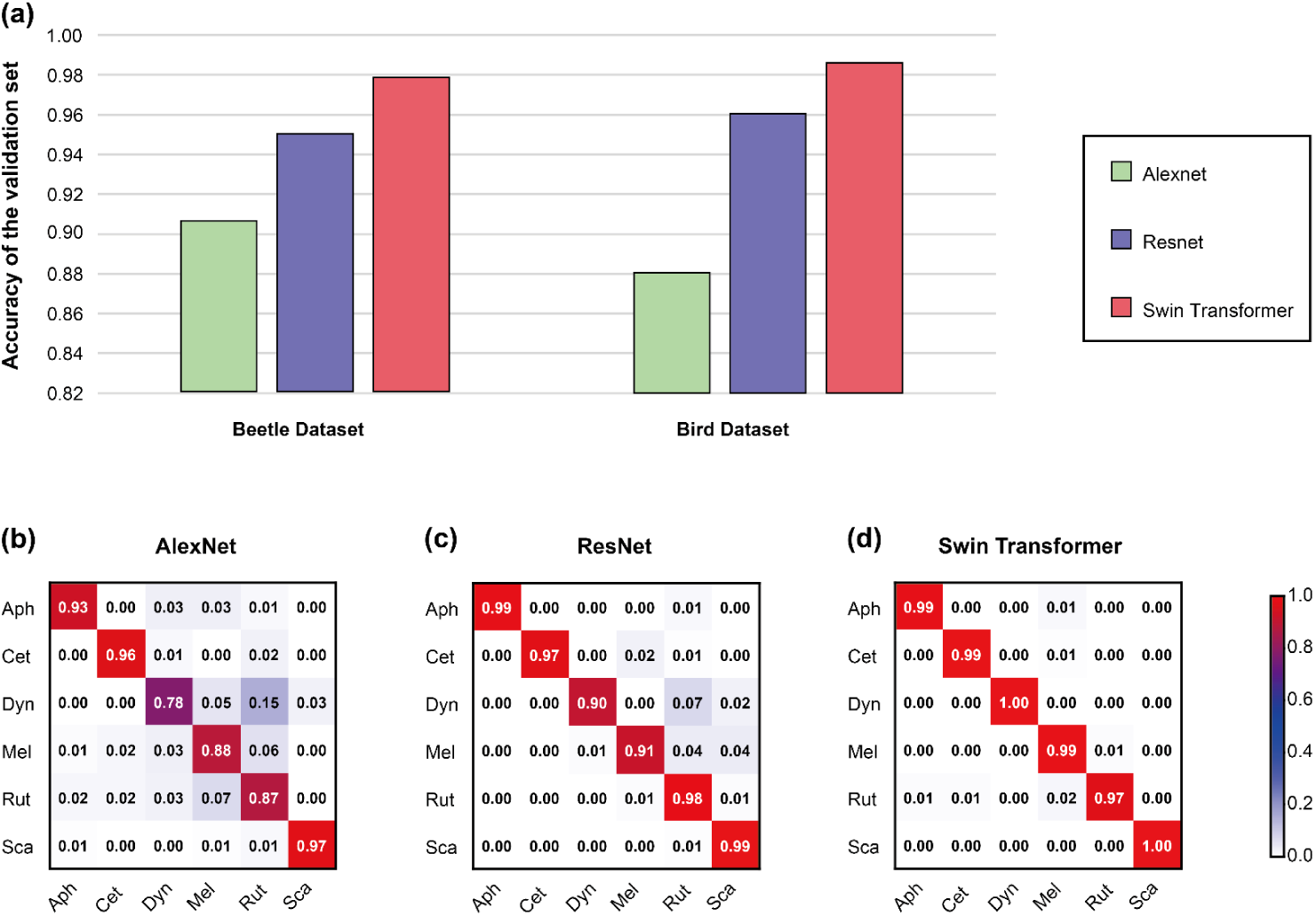
Comparison of the classification performance of AlexNet, ResNet, and Swin transformer. (a) The highest accuracy of the three models on the validation set. (b-d) Confusion matrix analysis of the three models on the beetle dataset.

#### 3.1.2 Bird dataset

On the bird dataset, AlexNet, ResNet, and Swin transformer also converged, with training times of 218, 487, and 478 minutes, respectively. The relatively simple network structure of AlexNet led to a shorter training time but a lower accuracy of only 88.5%, compared to ResNet’s 96.05% and Swin transformer’s 98.6% (Figure 2a). In confusion matrix analysis, Swin transformer still performs the best, with only 25 mispredicted samples out of 2000 (Figure S4). Therefore, on these two datasets, Swin transformer outperforms the other two models in both accuracy and confusion matrix analysis.

### 3.2 Retrieval capability of hash codes

Figure 3 presents the validation results for hash codes of varying lengths on six datasets, as evaluated by mAP. The highest mAP value on the beetle dataset occurs with a hash code length of 48, reaching 98.2%. For different datasets, the mAP values of hash codes retrieval are somewhat different. For the Fungi, Butterfly, Flower, Bird, and Fused datasets, the highest mAP values are achieved with hash code lengths of 128, reaching 88.9%, 99%, 95.4%, 97.6%, and 94.8%, respectively (Figure. 3a-e). Among them, we observed that the highest mAP value for the Fungi dataset with the complex backgrounds was 88.9%, indicating suggesting that the background of the dataset can have an impact on the retrieval accuracy of hash codes to some extent. Although the mAP value of the Fungi dataset is relatively low, its impact on the mAP value of the Fused dataset is weakened because mAP takes into account the average precision of all categories.

**FIGURE 3.**
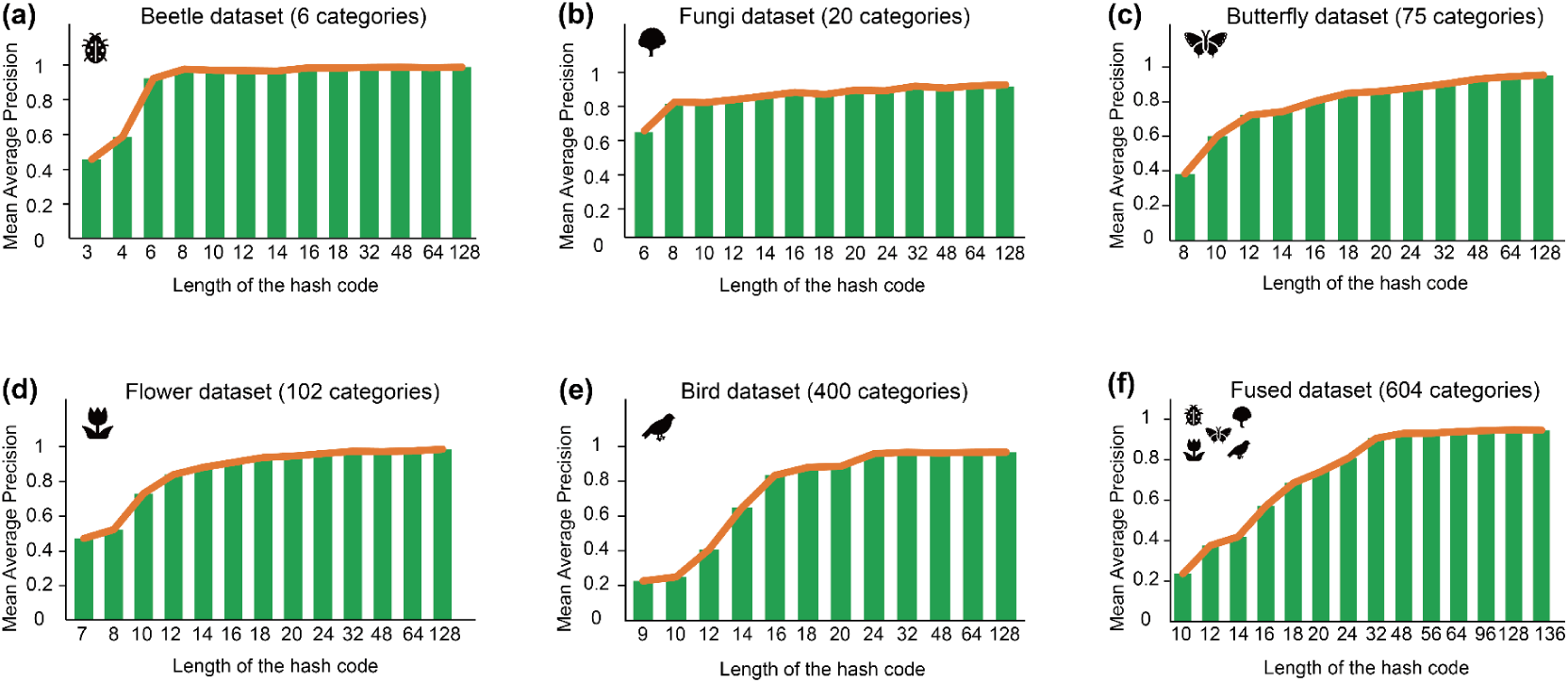
Histograms obtained under different hash code bits with their corresponding mAPs. a-e correspond to the beetle, fungi, butterfly, flower, and bird datasets, respectively, and f is fused from these five datasets. The number of categories in each dataset is parentheses after each heading.

Across all datasets, the mAP values exhibit a plateau pattern where they gradually increase as the length of hash codes increases, and then reach a stable level. This suggests that the length of the required hash code does not increase infinitely as the number of categories increases. In the six datasets analyzed in this study, it was found that once the hash code length exceeded 64 bits, the changes in mAP values became negligible. Therefore, in the simulated database scenario, we use 64-bit hash codes as a demonstration.

In the demonstration of the simulated database, after retrieving data from a simulated database using hash codes, the top five results all correspond to the same category as the hash code used for the retrieval (Figure S6). Since none of the images was involved in training, and no additional label information was provided during the retrieval process, this demonstrates that PENet extracted discriminative features between different categories. Even if we don’t have knowledge about the query image, the hash code can match individuals in the database with similar morphological features to the query image. Therefore, we can quickly retrieve the specimens based on morphological features once we generate a library of corresponding hash codes from existing digitized specimens. In this way, using hash codes for retrieval significantly improves efficiency and reduces storage space requirements. Taking the simulated database we built as an example, even if all the images are compressed to approximately 220×220 pixels, the storage space required for their images is still approximately 8000 times what is required for the hash codes.

### 3.3 Visualization of hash codes

After reducing the dimensionality of the hash code, which contains 64 feature values, using t-SNE, we can observe that groups of categories with similar features are still effectively clustered together (Figure S5a-e). This indicates that the hash codes carry sufficient discriminative features and provide a good representation of them, even on a bird dataset containing 400 categories, where intraclass and interclass distances are well distinguished. Furthermore, the visualization of discriminative features corresponding to each bit of the hash code also supports this view. For the 30 selected species, 64 heatmaps were generated for each, one of these species was displayed in Figure 4, and the rest of the results are detailed in supporting information. Here, the areas with the highest intensities in the heatmaps of status 1 in each bit of hash code cover the features extracted from the PENet, Status 0 means that these samples do not contain the features extracted in these hash code bits. After studying these heatmaps, it is further indicated that some bits of hash codes pointing to the real discriminative features which have been used in traditional ways, for example, bit 1: legs, bit 16: the shape of the prothorax, bit 39: the center of the body, and bit 44: the end of the abdomen.

**FIGURE 4.**
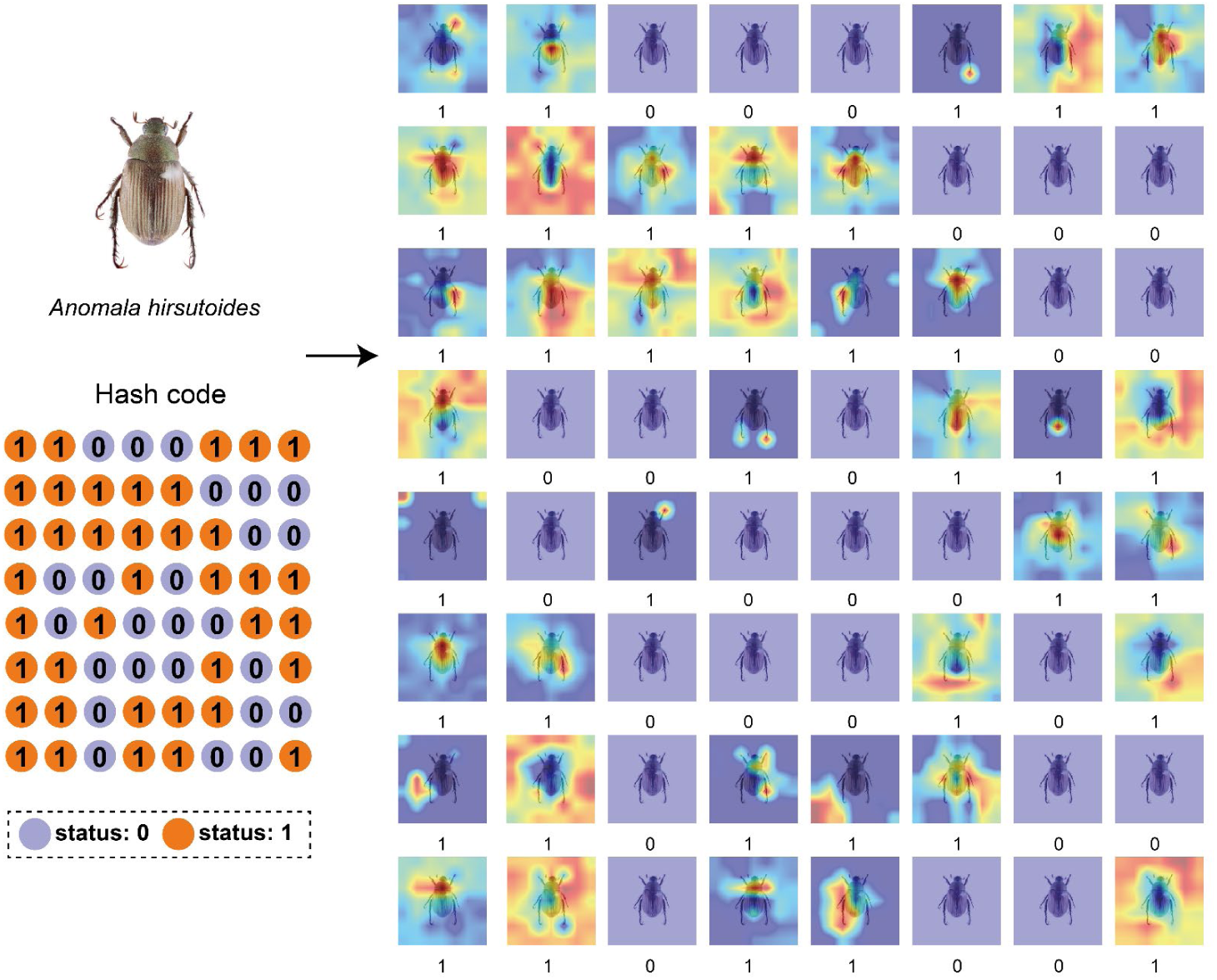
A 64-bit hash code is generated from a specimen by the PENet, where each bit corresponds to some features. The arrangement order of each bit in the hash code is from left to right, then from top to bottom.

### 3.4 Phenetic distance tree based on hash codes

We utilized PENet to convert 13 additional images of non-trained beetles into hash codes (Table S2). The hash codes of images belonging to the same category exhibited a similar pattern, whereas those of different categories showed some distinctions. For image that falls outside of the known categories (Staphylinidae), which can be considered out-of-distribution, its hash code differ from those of the known categories. To further investigate the value of hash codes in systematics, we constructed a phenetic distance tree. The maximum parsimony analysis of the 64 hash codes yields two most parsimonious trees (tree length=100 steps, CI=0.55, RI=0.72). “Morphological characters” (Hash codes) were optimized with parsimony on the first of the two most parsimonious trees (Figure 5), showing only “unambiguous” changes. Black circles indicate “nonhomoplasious” changes, and white circles indicate changes in “homoplasious characters”. The number above the branch represents “hash code” numbers, below branch are “hash code” status (0 or 1).

**FIGURE 5.**
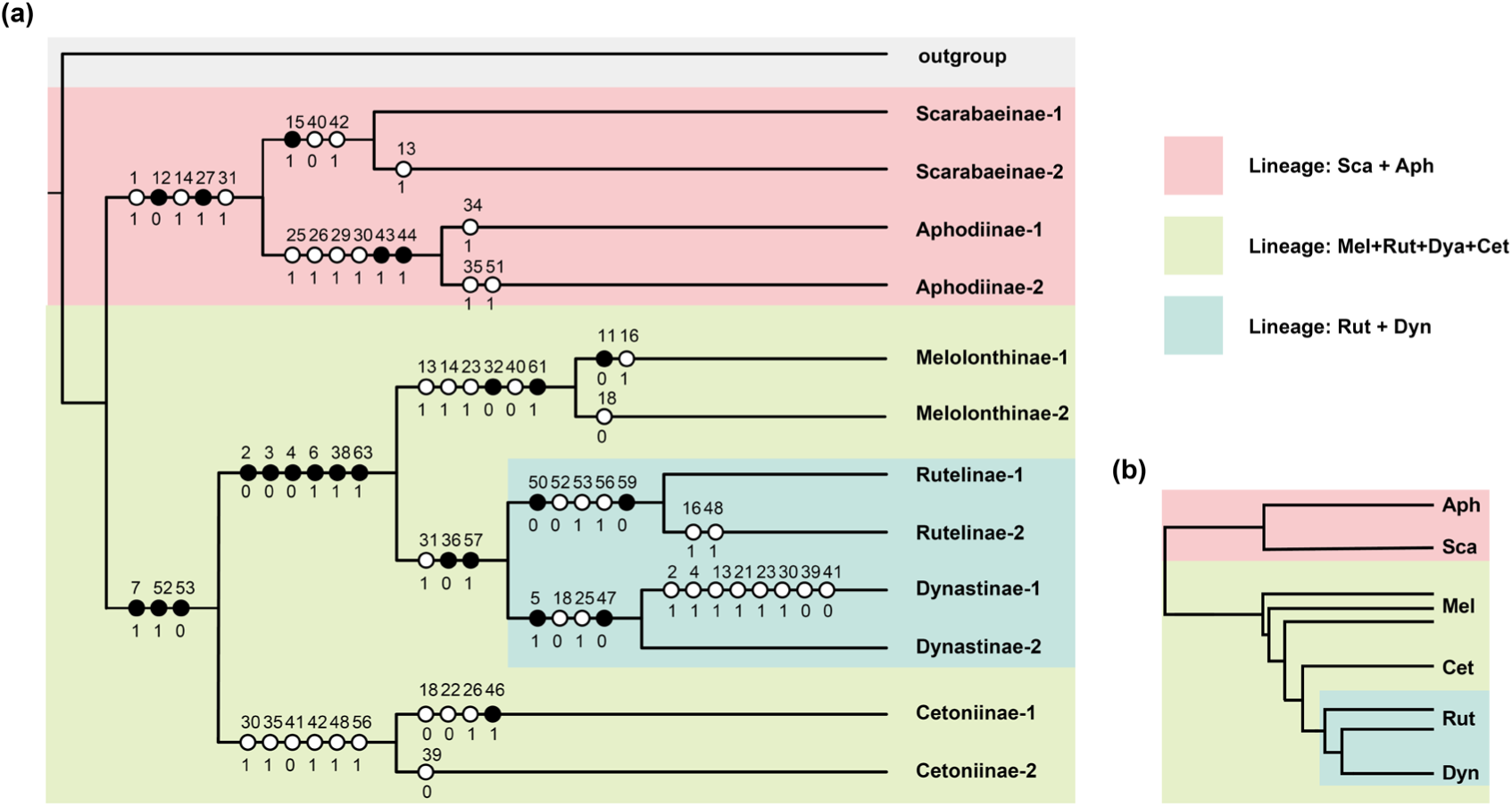
(a) Phenetic distance tree based on hash codes. Two species were selected for each subfamily with a hash code length of 64. The family Staphylinidae was selected as outgroup. (b) The simplified relationships among six subfamilies revealed by existing phylogenetic trees (Ahrens et al., 2014; Lu et al., 2023; Meinecke, 1975).

From the topological structure of the phenetic tree constructed by hash codes, it is shown that there are two basic lineages within the family of Scarabaeidae: coprophagous lineage (Scarabaeinae+Aphodiinae) and phytophagous lineage (Cetoniinae+Melolonthinae+Rutelinae+Dynastine), within phytophagous lineage Melolonthinae, Rutelinae, and Dynastine are cluster together, and the Dynastine and Rutelinae are sister groups. The topological structure of the phenetic tree constructed by hash codes similar to the phylogenetic trees revealed from molecular and morphological data, indicating that the hash codes contain additional distance information (Figure 5).

## Discussion

The implementation of automated and efficient discriminative feature extraction, along with mathematical encoding of extracted features, will further accelerate the development of research that relies on the dependence of morphological features. Here, we developed a novel deep learning model called PENet, which can rapidly extract discriminative features and represent them efficiently using hash codes. Our study has shown that the hash codes generated by PENet are an efficient carrier of the extracted discriminative features, as they encode these features as sequences. This encoding enables fast retrieval performance and facilitates comparisons and matches with minimal computational resources. In addition, we have also explored the potential application of hash codes in systematics and discovered their potential for further applications.

The Swin transformer, as the feature extractor of PENet, outperforms traditional convolutional neural networks in its ability to extract discriminative features. This can be seen in the confusion matrix analysis of the beetle dataset, where the Swin transformer performs well on each subfamily (Figure. 2d). In contrast, AlexNet and ResNet show imbalanced performance on each subfamily, with the main prediction errors occurring in Dynastinae and Melolonthinae. For Dynastinae, which had the smallest number of images, the imbalanced data distribution may have been a contributing factor to the poor performance of the AlexNet and ResNet models. For Melolonthinae, as the largest subfamily with numerous species (∼630 genera), there may be more extensive variations in the morphological features of its species, which could have impacted the performance of the models. Unlike AlexNet and ResNet, the Swin transformer excels in handling data imbalance and complex feature data. This may be attributed to its unique architecture, which segments images into small tokens to extract features and incorporates self-attention mechanisms to capture the interdependencies between different tokens, allowing for more effective feature extraction and weighting (Liu et al., 2021; Vaswani et al., 2017). Furthermore, some studies have shown that, unlike convolutional neural networks which focus on extracting texture information from images, transformer-based models place greater emphasis on extracting shape information from the global image (Baker et al., 2018; Tuli et al., 2021). Therefore, on the whole, Swin transformer has greater potential for extracting discriminative features, and it also endows PENet with improved performance.

The hash codes generated by PENet serve as carriers of the features extracted from the images and have been shown to possess strong representational power. The results of retrieval tests conducted on six datasets demonstrate that these hash codes are effective in representing discriminative information and enabling the retrieval of similar categories, even when the dimensionality of the data has been reduced. Furthermore, among the retrieval results of the same category based on the hash codes, the images that rank higher are more similar to the query image in terms of their features (Figure S6).

Natural history collections serve as a critical resource for studying the morphological traits of various species, and the digitization of these collections has significantly improved research in these fields (Lister, 2011; Page et al., 2015). Furthermore, with the application of machine learning and deep learning technologies, the utilization of these collections has been further optimized, leading to remarkable results in tasks such as rapid specimen identification (Younis et al., 2018), collections label information extraction (Owen et al., 2020), and functional traits measurement (Weeks et al., 2023). However, it is still challenging to quickly obtain specimens with similar morphology from large natural history collections. Our newly proposed deep learning model, PENet, can help address this issue. By using PENet to extract the morphological features of digital specimen images and converting these features into hash codes, we can quickly retrieve specimens with similar morphological characteristics. This approach also proves beneficial in cases where the taxonomic information of the specimens in question is uncertain. As hash codes are merely binary encodings, they do not add significant storage costs to digital collections. Furthermore, under ideal conditions, multiple natural history museums can share a set of hash codes as data accumulate.

Furthermore, the application of PENet is not limited to just natural history collections, but is suitable for any scenario that requires large-scale matching of morphological information, such as biodiversity monitoring. With the increasing deployment of infrared cameras in the wild, a vast amount of new data is generated every day (Burton et al., 2015). Relying solely on manual labor to search for target species in such a massive dataset can be challenging(Schneider et al., 2019). To address this, PENet can be used to train for target species, transforming all images into hash codes with feature information when searching for the target species. By comparing these hash codes with the target species’ hash codes, retrieval can be achieved in large-scale monitoring data. Additionally, with the aid of the corresponding algorithms, multiple ecological factors such as biodiversity and abundance can be obtained, promoting relevant ecological research.

In addition to enabling fast specimen retrieval, hash codes also have the potential to be further applied in systematics. Unlike most existing species classification models, PENet takes into account the distance relationship between different categories during the training process of generating hash codes. Therefore, the hash codes not only represent the extracted discriminative features but also carry distance information between different categories. Tests conducted on six subfamilies of the Scarabaeidae demonstrated that hash codes can be used to generate a phenetic distance tree. When compared with the existing phylogenetic tree, the phenetic distance tree showed some similarities in the division of certain major branches: two basic lineages, sister group relationship between Rutelinae and Dynastine (Ahrens et al., 2014; Lu et al., 2023; Mckenna et al., 2015). And the position of Cetoniinae is similar to some morphological-based phylogenetic results, which further indicate that the hash codes could reveal the phenetic distances and relationships among categories to a certain extent (Howden, 1982; Meinecke, 1975). However, our experiment was only preliminary as we only used the dorsal view, which was not sufficient to cover all features of the test species. In future research, we will continue to supplement the dataset and use methods such as multi-angle photography and microscopic photography to obtain comprehensive morphological information and further validate the ability of the hash codes.

There are still some issues to consider to better implement PENet in practice. Since the length of the hash code and the specificity of the training data affect the achieved retrieval performance, the optimal hash code length should be determined according to the actual needs of different fields and the training data should be carefully selected to minimize the influence of the hash code length on the performance of the PENet. Additionally, although the features extracted by PENet are similar to those perceived by experts to a large extent, additional confirmation should be required to confirm whether the extracted features match the phenotypic information to be studied.

## Conclusion

Overall, our newly developed end-to-end PENet model demonstrates excellent performance in feature extraction and fast retrieval, with the potential for broader applications in systematics. Hash codes carry both discriminative features and phenetic distance information while maintaining a low-dimensional representation, allowing efficient morphological information retrieval with a minimal storage cost. PENet provides an effective solution for the fast retrieval of natural history collections. Furthermore, it can be further applied to explore morphological features, supporting research on macro-morphological evolution and mimicry. In future research, we will continue to explore the extended applications of hash codes and consider unsupervised training methods that rely solely on morphological distance information to train PENet, further increasing its scope of applications.

## ACKNOWLEDGMENTS

We thank Jing Li and Ning Liu from the Institute of Zoology, Chinese Academy of Sciences for technical support in computers. And we thank Chuanbu Gao for providing some beetle images for this study. Shiqing Qiao, Yiyao Zhang, and Zi Jin from Shenyang Normal University, and Jiateng Zhao from Hebei University for data collecting. This research was funded by the National Key R&D Program of China (Grant No. 2022YFC2601200); the National Natural Science Foundation of China (No. 32270468, 32200354, 31961143002); and China Postdoctoral Science Foundation (No. 2022M713134).

## AUTHOR CONTRIBUTIONS

Z. Y. Z., Y.Y. L., and M. B. designed the study and wrote the paper. Z. Y. Z. performed the training of the model, and Y.Y. L. interpreted the results. Z. Y. Z., Y. J. T., and X.C. completed the data collection. X.C. managed the equipment for model training. All authors read and approved the manuscript.

---

**SUPPLEMENTARY TABLE 1.**
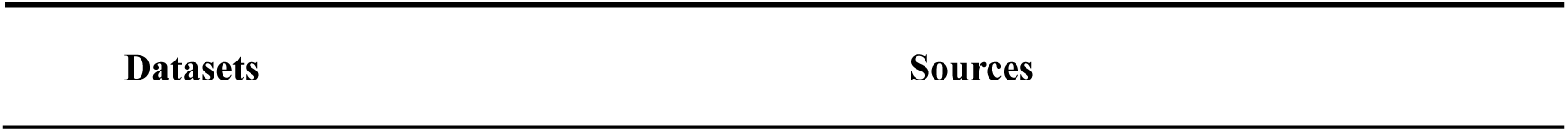
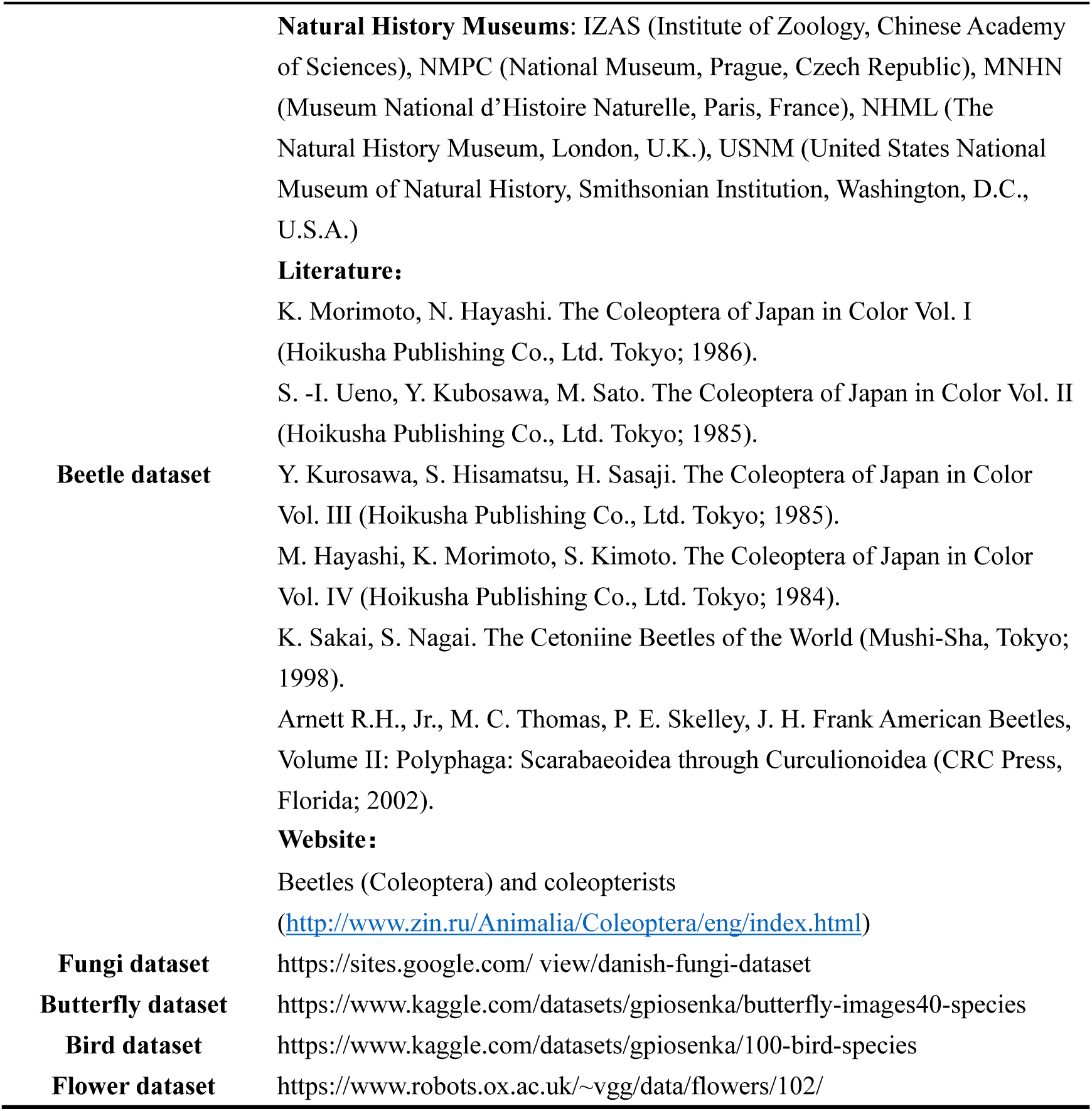
Sources of the datasets.

**SUPPLEMENTARY TABLE 2.**
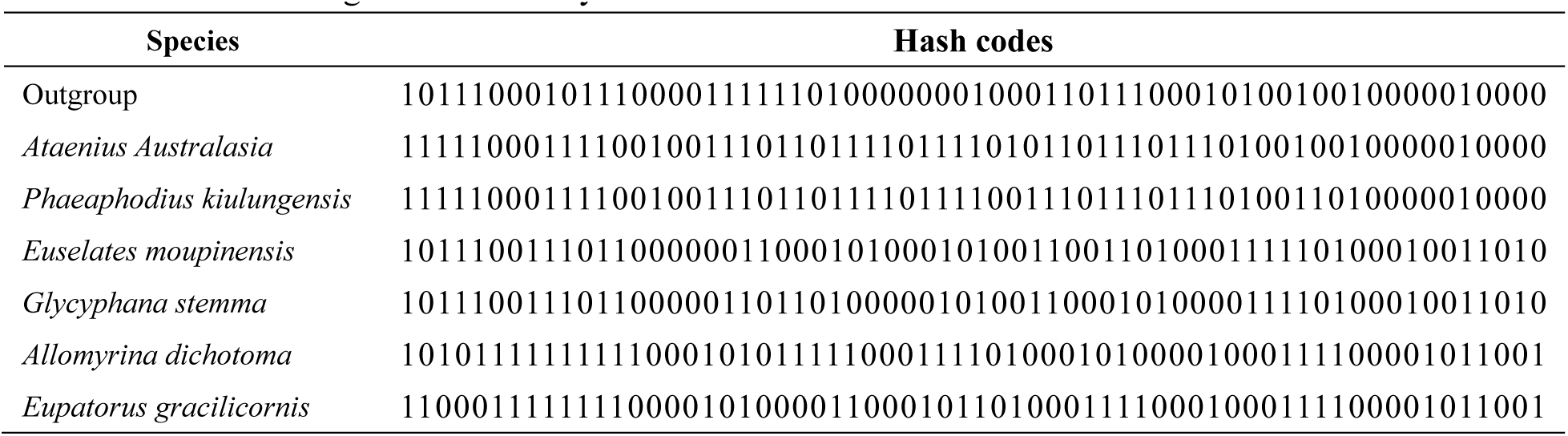
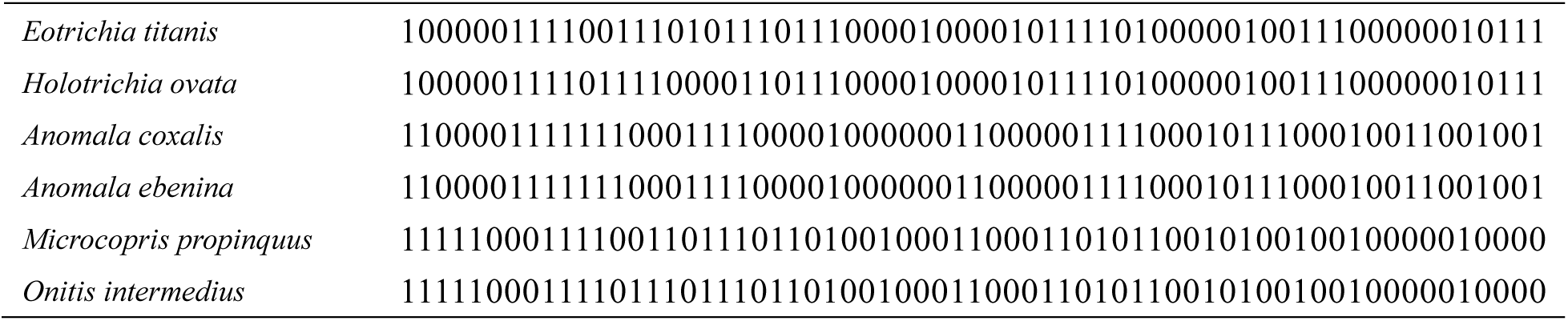
The species used to build the phenetic tree and the corresponding hash codes. Among them, *Ataenius australasiae* and *Phaeaphodius kiulungensis* belong to the subfamily Aphodiinae, *Euselates moupinensis* and *Glycyphana stemma* belong to the subfamily Cetoniinae, *Allomyrina dichotoma* and *Eupatorus gracilicornis* belong to the subfamily Dynastinae, *Eotrichia titanis* and *Holotrichia ovata* belong to the subfamily Melolonthinae, *Anomala coxalis* and *Anomala ebenina* belong to the subfamily Rutelinae, *Microcopris propinquus* and *Onitis intermedius* belong to the subfamily Scarabaeinae.

**SUPPLEMENTARY FIGURE 1.**
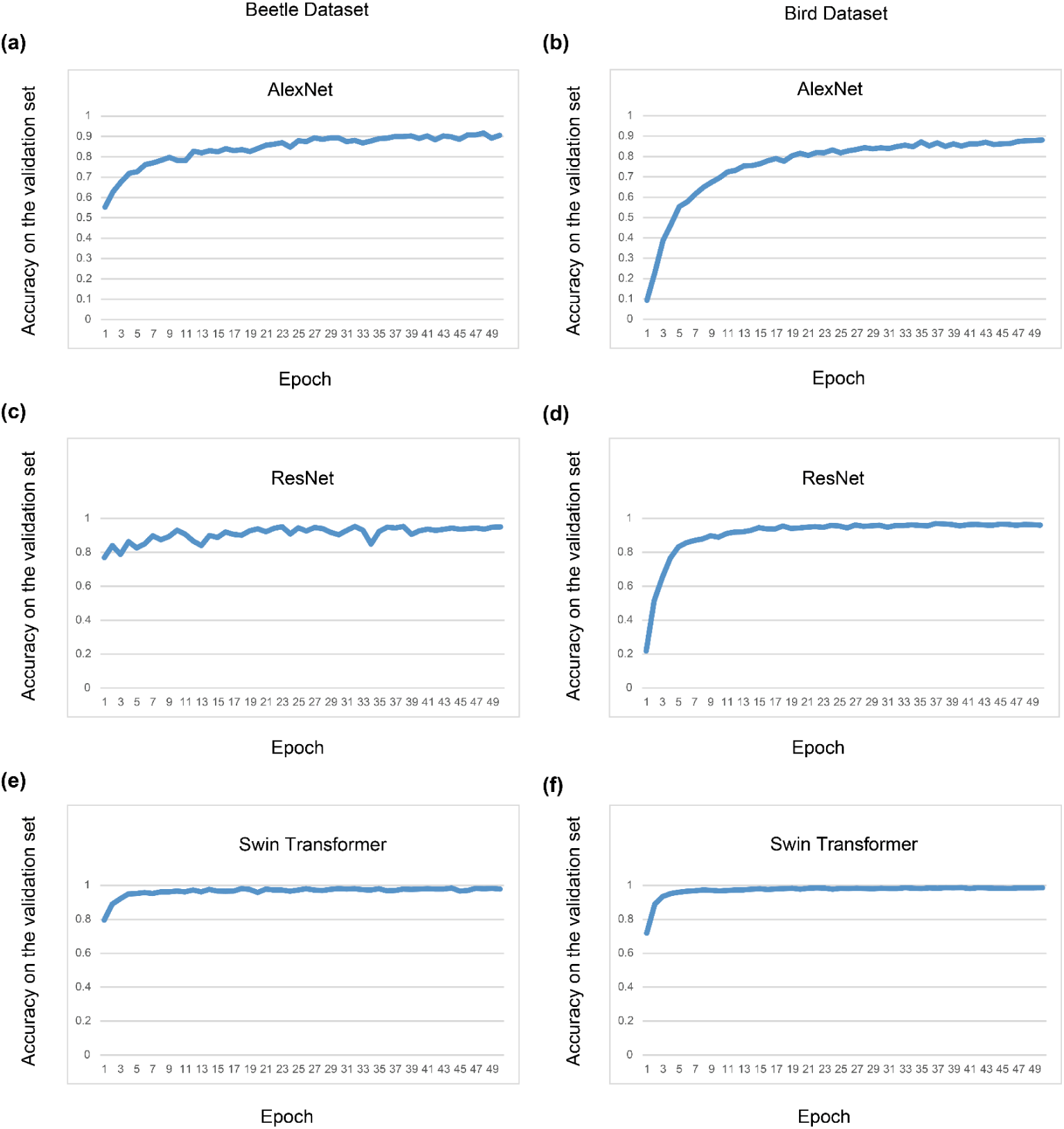
Classification accuracy of AlexNet, ResNet, and Swin transformer in the validation set of beetles and birds.

**SUPPLEMENTARY FIGURE 2.**
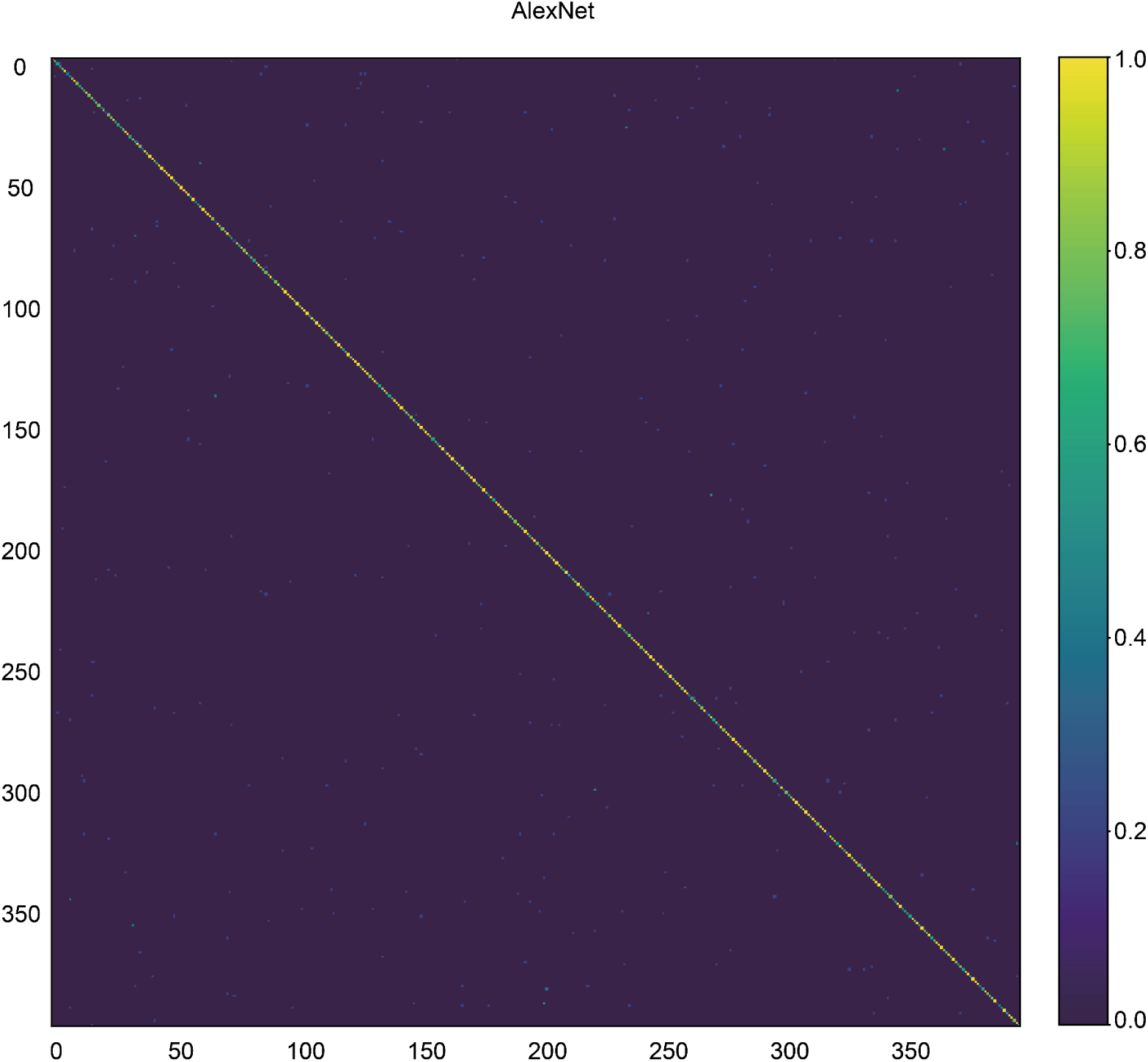
Confusion matrix analysis of AlexNet on the bird dataset.

**SUPPLEMENTARY FIGURE 3.**
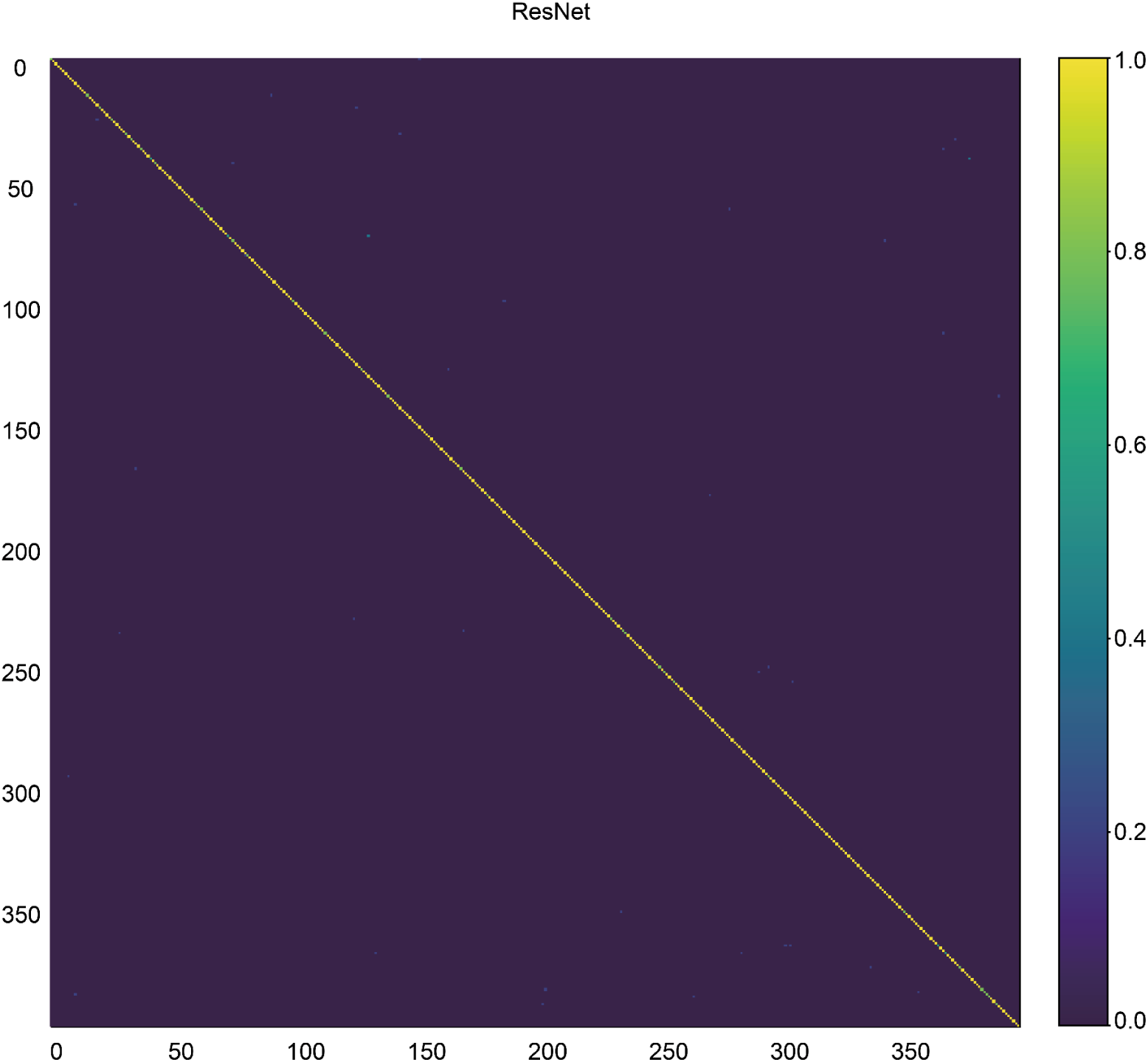
Confusion matrix analysis of ResNet on the bird dataset.

**SUPPLEMENTARY FIGURE 4.**
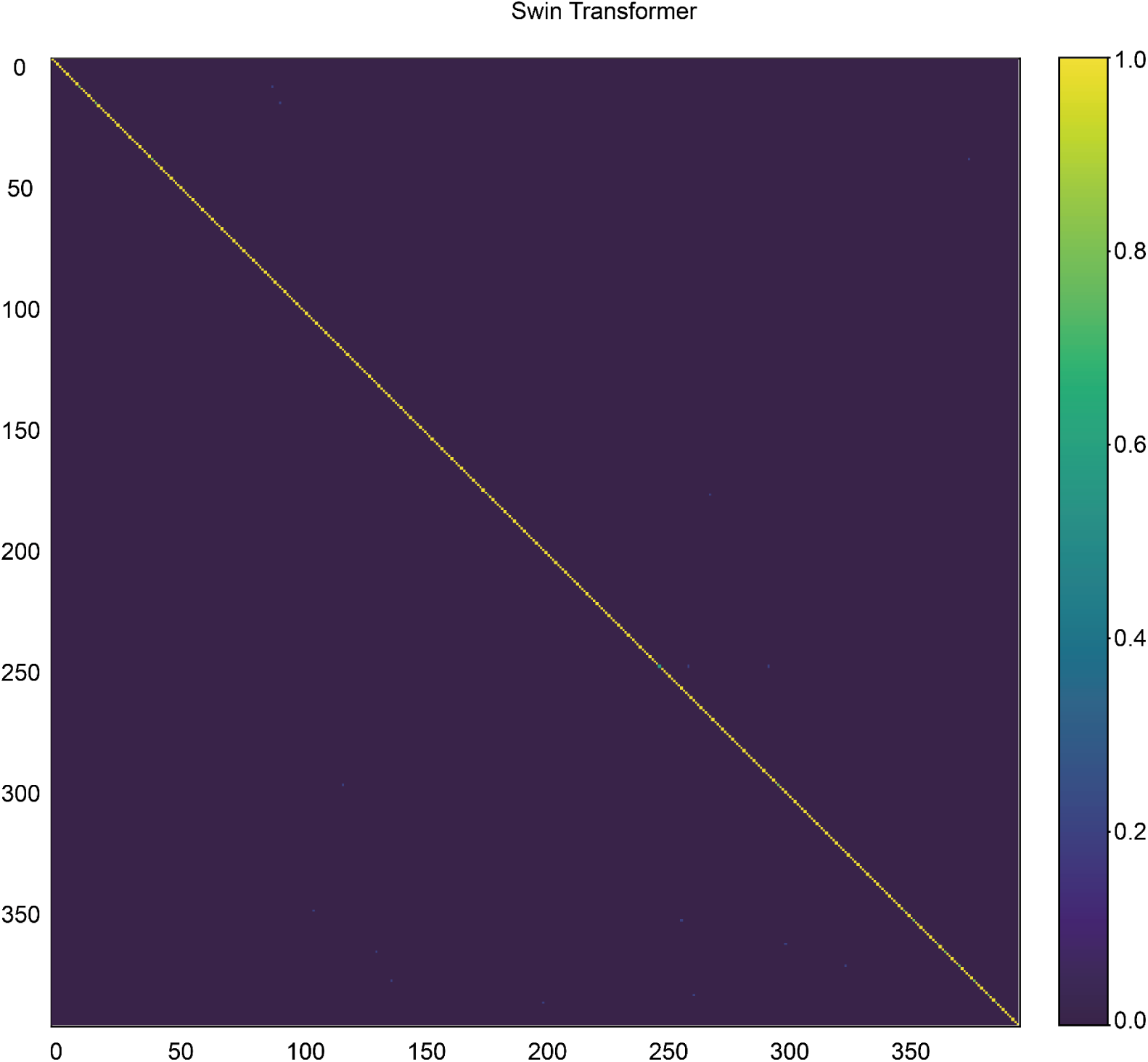
Confusion matrix analysis of Swin transformer on the bird dataset.

**SUPPLEMENTARY FIGURE 5.**
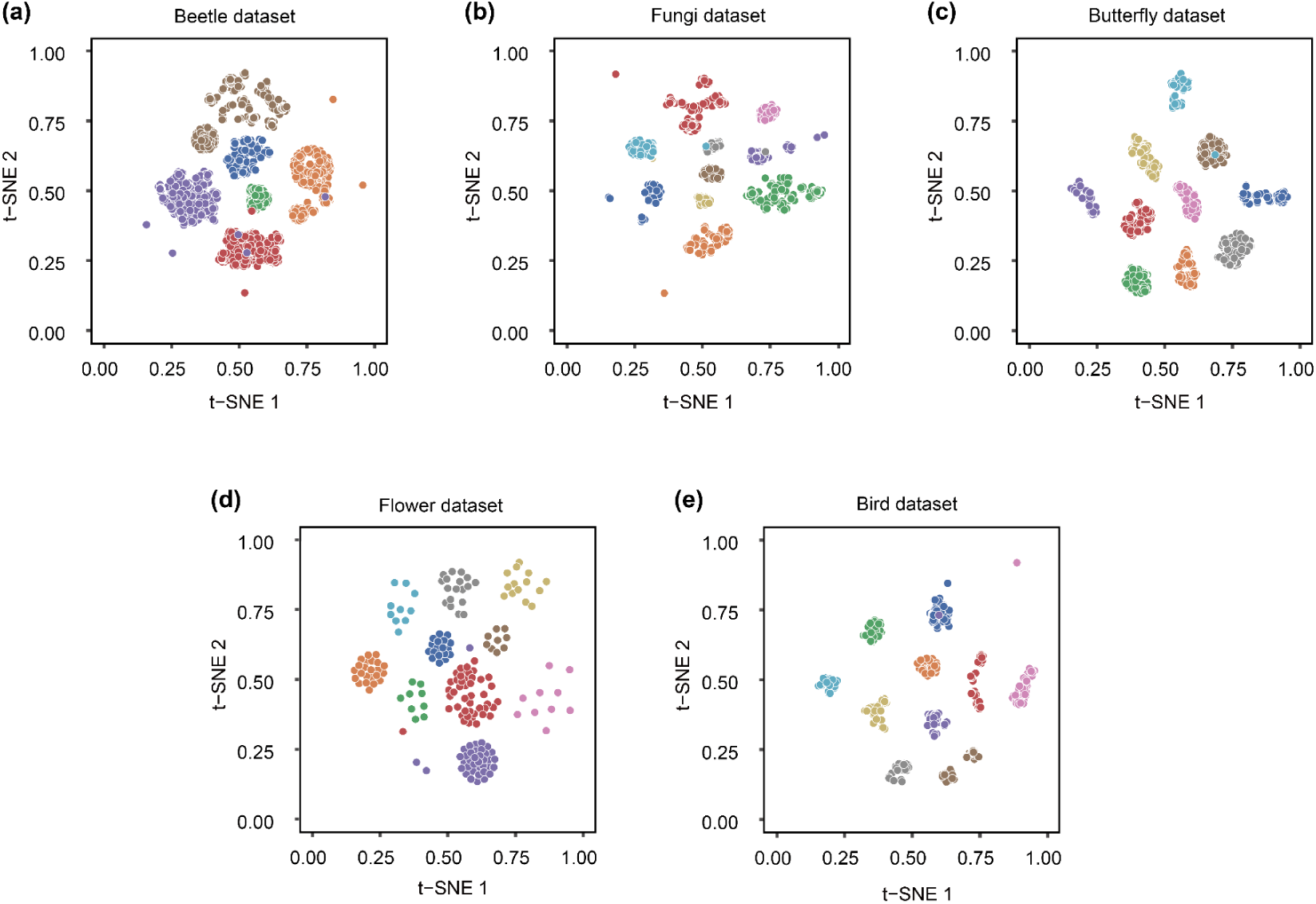
Scatter plots of 64-bit hash codes as feature data after performing t-SNE dimensionality reduction, where (a-e) are obtained by randomly drawing 10 classes from the corresponding dataset.

**SUPPLEMENTARY FIGURE 6.**
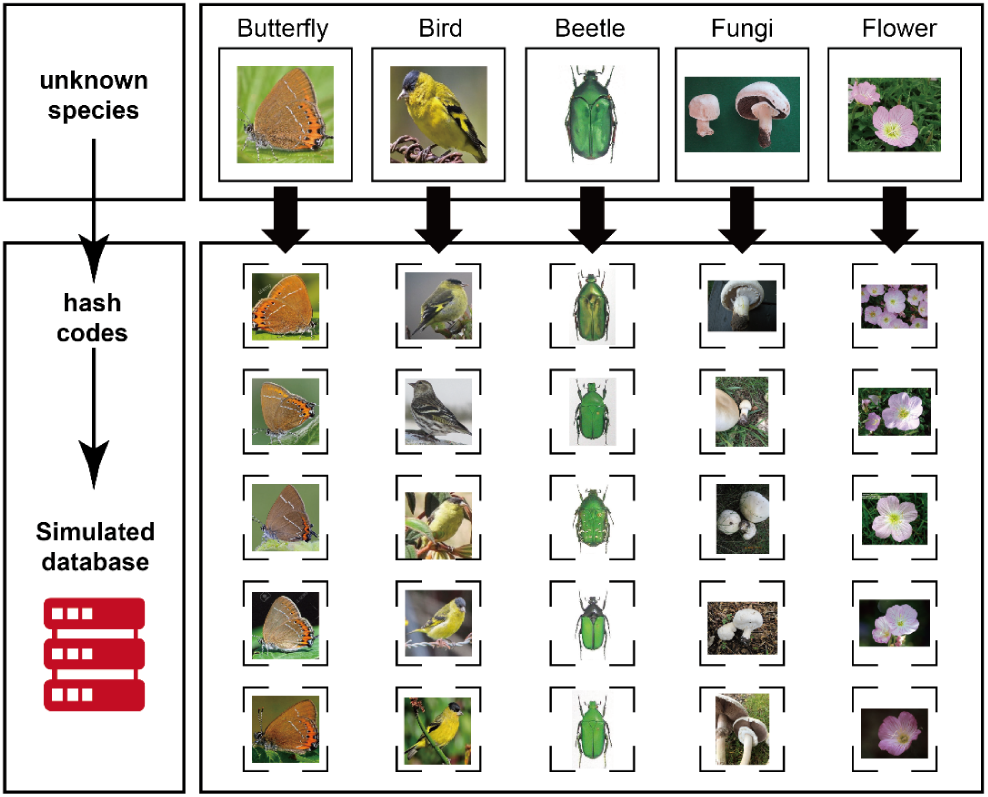
Retrieval results with hash codes as indexes. For each data set, one image is randomly selected and used as an index to retrieve the top five images with the highest similarity to it in the simulated database.

